# Association between peripheral inflammation and DATSCAN data of the striatal nuclei in different motor subtypes of Parkinson Disease

**DOI:** 10.1101/247080

**Authors:** Hossein Sanjari Moghaddam, Farzaneh Ghazi Sherbaf, Mahtab Mojtahed Zadeh, Amir Ashraf-Ganjouei, Mohammad Hadi Aarabi

## Abstract

The interplay between peripheral and central inflammation has a significant role in dopaminergic neural death in nigrostriatal pathway, although no direct assessment of inflammation has been performed in relation to dopaminergic neuronal loss in striatal nuclei. In this study, the correlation of neutrophil to lymphocyte ratio (NLR) as a marker of peripheral inflammation to striatal binding ratios (SBR) of DAT-SPECT images in bilateral caudate and putamen nuclei were calculated in 388 drug-naïve early PD patients (288 tremor-dominant, 73 PIGD, and 27 intermediate) and 148 controls. NLR was significantly higher in PD patients than age and sex-matched healthy controls. NLR showed a negative correlation to SBR in bilateral putamen in all PD subjects. Among our three subgroups, only TD subgroup showed remarkable results. A positive association between NLR and motor severity was observed in TD subgroup. Besides, NLR could negatively predict the SBR in ipsilateral and contralateral putamen and caudate nuclei in tremulous phenotype. Nonetheless, we found no significant association between NLR and other clinical and imaging findings in PIGD and intermediate subgroups, supporting the presence of distinct underlying pathologic mechanisms between tremor and non-tremor predominant PD at early stages of the disease.

## Introduction

Progressive dopaminergic demise in the nigrostriatal system has long been considered the pathogenic mainstay of motor features in Parkinson’s disease (PD). Lots of ergot mechanisms have been enumerated responsible for the neural loss in PD, of which inflammation, both central and peripheral has been explained in the initiation and progression of PD.

Inflammatory processes, which once were believed to be just the afterclap of neural death, are now considered as an etiological factor, which are also presented in early stages of PD. Several PD susceptibility genes are documented to be involved in the functions of immunity components like microglia (Liu et al., 2011; Pihlstrom et al., 2013; Russo et al., 2014). The result from polymorphism studies determined several pro-inflammatory genes such as IL-1β, TNF-α, HLA-DBQ1, HLA-DRA, and HLA-DRB1 to be associated with increased susceptibility to PD (Wahner et al., 2007; Ahmed et al., 2012). It is also proved that nonsteroidal anti-inflammatory (NSAIDs) can in long term diminish the risk of PD (Rees et al., 2011). In essence, authors exhibited that dopaminergic neurons are selectively susceptible to oxidative stress and inflammation (Pott Godoy et al., 2008; Liu and Bing, 2011). Freshly, a neuroimaging study using positron emission tomography (PET), detected activated microglia in the SN and putamen of one-year duration PD patients (Iannaccone et al., 2013), implying that there is an active inflammatory condition in initial phases of PD. Furthermore, authors suggested that inflammatory circumstances, both central and peripheral, are present in the prodromal PD (Su and Federoff, 2014; Dzamko et al., 2015). Altogether, these immense lines of evidence highly signify the importance of inflammation in the pathophysiology of PD.

A robust body of evidence has espoused the involvement of peripheral inflammatory conditions in the pathophysiology of PD. In this context, markers of peripheral inflammatory processes including, but not limited to, TNF-α and its receptors and IL-1β have been exhibited to be elevated in in individuals with PD in the peripheral blood (Dobbs et al., 1999; Katsarou et al., 2007; Scalzo et al., 2009; Koziorowski et al., 2012; Rocha et al., 2014), cerebrospinal fluid and post-mortem brain tissues (Mogi et al., 1994; Dobbs et al., 1999; Reale et al., 2009a; Reale et al., 2009b). Related data showed IL-6 is associated with increased risk of development of PD (Scalzo et al., 2010). Besides the peripheral immune markers, peripheral immune cells undergo some quantitative alternations in PD patients. The most prominent of these changes is the lower total number of lymphocytes in individuals with PD compared to controls mainly due to decrement in percentage of CD3+ and CD19+ lymphocytes (Bas et al., 2001; Baba et al., 2005; Saunders et al., 2012; Stevens et al., 2012). Furthermore, decreased counts of T helper cells and increased/unchanged counts of cytotoxic T cells have been documented in PD patients (Calopa et al., 2010).

Owing to the presence of blood-brain-barrier (BBB), central nervous system (CNS) is an immune-privileged organ. However, BBB dysfunction have been postulated to play a key role in the pathogenesis of neurodegenerative diseases like PD (Kortekaas et al., 2005; Hirsch and Hunot, 2009). It has been shown that endothelium residing in SNpc of patients with PD, undergoes radical pathologic morphological changes (Faucheux et al., 1999; Guan et al., 2013), and CD4+ and CD8+ T-lymphocytes have been reported to enter and invade the SNpc of idiopathic PD patients (Phani et al., 2012). Using a mouse model involving injection of vascular endothelial growth factor (VEGF) (a detrimental protein to BBB) into the mice SNpc, researchers demonstrated a significant association between BBB dysfunction and dopaminergic cell death (Rite et al., 2007; Yasuda et al., 2007)., Furthermore, inflammatory cytokines can induce the activation of microglia and therefore the dopaminergic cell death. In this context, one of the valid indicators for peripheral inflammatory processes is the neutrophil to lymphocyte ratio (NLR) (Alkhouri et al., 2012) representing the active inflammation (neutrophils) merged with immunological regulatory processes (lymphocytes).

Despite lots of evidence on the role of inflammation in dopaminergic neural loss, no direct assessment has been performed in this regard. This work was designed to investigate peripheral inflammation in terms of NLR concerning loss of dopaminergic uptake in caudate and putamen on Dopamine Transporter Single Photon Emission Computed Tomography (DaTscan) images, which signifies the dopaminergic activity, among treatment-naïve early PD patients with different motor phenotypes and severities.

## 2. Methods

### 2.1 Participants

PD patients and healthy control (HC) subjects enrolled in this study were recruited from the Parkinson Progression Markers Initiative (PPMI, http://www.ppmi-info.org/). The study was approved by the institutional review board of all participating sites. Written informed consent was obtained from all participants before study enrolment. The study was performed in accordance with relevant guidelines and regulations. Only baseline visit data are analyzed in this research. PD status was confirmed by Movement Disorder Society-Unified Parkinson’s Disease Rating Scale (MDS-UPDRS) and dopamine transporter deficit was observed on DAT scans. All PD patients at baseline were drug naïve, non-demented, on Hoehn and Yahr (H&Y) staging I or II, and were confirmed negative for any medical or psychiatric disorders apart from PD.

### 2.2 Clinical assessment and motor classification

Motor and non-motor symptoms were evaluated among subjects with clinical tests at baseline visits at each participating site. The most common PD rating scales, i.e., UPDRS, H&Y staging, and the Schwab and England rating of activities of daily living (Modified Schwab & England ADL) were assessed (Perlmutter, 2009). Tremor score and postural instability and gait difficulty (PIGD) score were identified for PD subjects with MDS-UPDRS. Then PD subjects were classified into 3 groups of tremor dominant (TD), PIGD and intermediate based on the ratio of tremor score/PIGD score. If ratio >= 1.15 or PIGD=0 and tremor>0, then the subject was TD. If ratio <= 0.9, the subject was classified as PIGD. If ratio > 0.9 and < 1.15, or tremor score and PIGD score = 0, the subject was tagged as indeterminate (Stebbins et al., 2013). Non-motor symptoms were investigated by the Montreal Clinic Assessment (MoCA) for mild cognitive impairment, 15-item geriatric depression scale (GDS) for depressive symptoms, The University of Pennsylvania Smell Identification Test (UPSIT) for olfaction function, and independent validation of the scales for outcomes in Parkinson’s disease-autonomic (SCOPA-AUT) for autonomic dysfunction.

### 2.3 Blood sample collection

As it is pronounced in the PPMI biologics manual (http://www.ppmi-info.org/), baseline blood sample collection was completed at each study site. PAXgene blood RNA tubes were utilized for the collection of blood samples following the study protocol. The number of neutrophils and lymphocytes were calculated by autoanalyzer device on participants’ whole blood samples, right after the collection. NLR was simply calculated by dividing neutrophil to lymphocyte count.

### 2.4 Dopaminergic imaging

As a preventive measure, all females with the potential of pregnancy were confirmed negative for urine pregnancy test prior to receiving the ^123^Ioflupane single-photon emission computed tomography (SPECT) injection. None of the subjects had received any of the drugs that would interfere with DAT SPECT imaging within 6 months of screening (full PPMI study protocol is provided at http://www.ppmi-info.org/study-design/research-documents-and-sops/). DAT imaging at baseline visit from all participating cites were centrally reconstructed and were attenuation corrected, followed by spatial normalization for consistent orientation by experienced nuclear medicine experts. Standard volume of interest template was applied on caudate, putamen and occipital regions. Striatal Binding Ratios (SBR) for left and right putamen and caudate were finally calculated, using the formula: (SBR) = (striatal region)/(occipital) -1, where occipital lobe DAT count is the reference (Seibyl et al., 2013). Regarding motor dominancy, SBRs of ipsilateral and contralateral putamen and caudate were compared between PD and HC and within PD motor subtypes. The calculated average of SBRs for the left and the right nuclei is presented for healthy controls or symmetrical motor involvement in PD individuals.

### 2.5 Statistical analysis

The statistical analysis was performed using SPSS version 22 (BM Corp., Armonk, NY, USA). Pearson’s chi-square was used to assess nominal variables across groups. Mann-Whitney U test was used to assess differences between healthy controls and PD patients, and Kruskal-Wallis test was used for multiple comparisons in three groups of PD motor subtypes. Spearman’s rank-order correlation was used to test the association between NLR and other characteristics in patients with Parkinson’s disease. Finally, *P* values less than 0.05 were considered to be statistically significant.

## 3. Results

### 3.1 Between group comparisons

NLR and DAT scan data were provided for 388 de-novo PD patients and 148 HC on baseline visits. Baseline clinical characteristics and detailed demographic data of PD and HC subjects are demonstrated in table 1. Healthy participants were age and sex matched to PD patients, while scored higher on all motor and non-motor tests. As expected, PD patients had significantly lower SBRs in striatal nuclei. NLR was significantly higher in PD cohort.

**Table 1.** Demographic information and comparison of clinical outcomes between healthy controls and patients with Parkinson’s disease.

| Characteristic | HCs<br>(n = 148) | PD<br>(n = 388) | P Value |
| --- | --- | --- | --- |
| Age, mean (SD) [95% CI], y | 61.1 (11.3) [59.3-62.9] | 61.7 (9.6) [60.7-62.6] | 0.775<sup>a</sup> |
| Female/male, No. (% male) | 49/99 (66.9) | 134/254 (65.4) | 0.755<sup>b</sup> |
| Left-handed/right-handed, No.<br>(%right-handed) × | 22/119 (80.4) | 33/346 (89.2) | 0.029<sup>b</sup> |
| Education, mean (SD) [95% CI], y | 16.1 (3.0) [15.7-16.6] | 15.5 (3.0) [15.2-15.8] | 0.034<sup>a</sup> |
| Disease duration, median (range), m | ... | 4.0 [0-36] | ... |
| NLR, mean (SD) [95% CI] | 2.2 (0.8) [2.1-2.3] | 2.5 (0.9) [2.4-2.6] | <0.001<sup>a</sup> |
| SBR score in Caudate nucleus, mean<br>(SD) [95% CI] | 3.0 (0.6) [2.9-3.1] | 1.8 (0.6) [1.8-1.9] | <0.001<sup>a</sup> |
| SBR score in Putamen nucleus mean<br>(SD) [95% CI] | 2.1 (0.5) [2.0-2.2] | 0.7 (0.3) [0.6-0.7] | <0.001<sup>a</sup> |
| Hoehn and Yahr stage, mean (SD) | ... | 1.6 (0.5) | ... |
| MDS-UPDRS part I score,<br>mean (SD) | 3.0 (2.7) | 5.6 (4.0) | <0.001<sup>a</sup> |
| MDS-UPDRS part II score,<br>mean (SD) | 0.4 (1.0) | 6.0 (4.2) | <0.001<sup>a</sup> |
| MDS-UPDRS part III score,<br>mean (SD) | 1.2 (2.1) | 20.4 (8.8) | <0.001<sup>a</sup> |
| Modified Schwab & England ADL,<br>mean (SD) | ... | 93.2 (6.0) | ... |
| MoCA score, mean (SD) | 28.2 (1.1) | 27.1 (2.3) | <0.001<sup>a</sup> |
| GDS score, mean (SD) | 1.1 (1.5) | 2.3 (2.4) | <0.001<sup>a</sup> |
| UPSIT score, mean (SD) | 34.4 (4.5) | 22.3 (8.2) | <0.001<sup>a</sup> |
| SCOPA-AUT score, mean (SD) | 5.7 (3.5) | 9.4 (6.0) | <0.001<sup>a</sup> |

Abbreviations: HCs, healthy controls; PD, Parkinson disease; NLR, Neutrophil to lymphocyte ratio; SBR, Striatal binding ratios; MDS-UPDRS, Movement Disorder Society-sponsored revision of the Unified Parkinson's Disease Rating Scale; MoCA, Montreal cognitive assessment; GDS, Geriatric depression scale; Modified Schwab & England ADL, overall activities of daily living; UPSIT, University of Pennsylvania Smell Identification Test; SCOPA-AUT, Scales for Outcomes in Parkinson's disease.
× Others were mixed-handed
<sup>a</sup>Based on Mann-Whitney *U* test.
<sup>b</sup>Based on $\chi^2$ test.

As shown in table 2, there was no difference between age, gender, UPDRS-III as well as NLR and SBRs in both ipsilateral and contralateral striatal nuclei among PD motor subtypes.

**Table 2.** Demographic data and comparison of certain characteristics between patients with Parkinson’s disease based on Tremor/PIGD score ratio.

| Characteristic |  | PIGD *<br>(n = 73) | Indeterminate<br>(n = 27) | TD<br>(n = 288) | P Value |
| --- | --- | --- | --- | --- | --- |
| Age, mean (SD) [95% CI],<br>y |  | 60.8 (9.5)<br>[58.6-63.0] | 62.8 (10.0)<br>[58.8-66.8] | 61.8 (9.7)<br>[60.7-62.9] | 0.744 <sup>a</sup> |
| Female/male, No. (% male) |  | 29/44<br>(60.0) | 6/21 (77.7) | 99/189 (65.6) | 0.261 <sup>b</sup> |
| MDS-UPDRS part III<br>score,<br>mean (SD) [95% CI] |  | 19.2 (8.7)<br>[17.1-21.2] | 18.1 (7.7)<br>[15.1-21.2] | 20.4 (9.0)<br>[19.9-22.0] | 0.195 <sup>a</sup> |
| NLR, mean (SD) [95% CI] |  | 2.5 (0.8)<br>[2.3-2.6] | 2.6 (1.0)<br>[2.2-3.0] | 2.5 (1.0)<br>[2.4-2.6] | 0.952 <sup>a</sup> |
| SBR in<br>contralateral<sup>^</sup><br>nuclei, mean<br>(SD) [95% CI] | Caudate | 1.8 (0.6)<br>[1.6-1.9] | 1.8 (0.5)<br>[1.6-2.0] | 1.8 (0.6)<br>[1.8-1.9] | 0.631 <sup>a</sup> |
|  | Putamen | 0.7 (0.3)<br>[0.6-0.8] | 0.6 (0.2)<br>[0.5-0.7] | 0.7 (0.3)<br>[0.6-0.7] | 0.163 <sup>a</sup> |
| SBR in<br>ipsilateral<br>nuclei, mean<br>(SD) [95% CI] | Caudate | 2.1 (0.7)<br>[2.0-2.3] | 2.1 (0.6)<br>[1.9-2.4] | 2.2 (0.6)<br>[2.1-2.2] | 0.813 <sup>a</sup> |
|  | Putamen | 1.0 (0.4)<br>[0.8-1.0] | 1.0 (0.4)<br>[0.8-1.2] | 1.0 (0.4)<br>[0.9-1.0] | 0.713 <sup>a</sup> |

Abbreviations: NLR, Neutrophil to lymphocyte ratio; SBR, Striatal binding ratios; MDS-UPDRS, Movement Disorder Society-sponsored revision of the Unified Parkinson's Disease Rating Scale.
\*The ratio of tremor and PiGD score in MDS-UPDRS. Parkinson's disease patients with a ratio of $\geq 1.15$ were classified as tremor-dominant (TD), ratio $\leq 0.90$ were classified as PiGD and $0.90 < \text{ratio} < 1.15$ were classified as intermediate.
<sup>^</sup>Contralateral and ipsilateral striatal nuclei are against the side of motor dominance.
<sup>a</sup>Based on Kruskal-Wallis test.
<sup>b</sup>Based on $\chi^2$ test.

### 3.2 Correlations between NLR and other clinical and imaging metrics

Spearman’s rank-order correlation was performed in healthy controls and three subgroups of PD patients (Table 3). NLR had a strong positive correlation with age and UPDRS III score and a negative correlation with Modified Schwab & England ADL score only in TD subgroup of PD patients. Moreover, TD patients showed negative correlation between NLR values and SBR in contralateral putamen (p-value 0.041) and ipsilateral putamen and caudate regarding motor dominancy (P-values <0.001 and 0.005 respectively). No significant correlation was observed between NLR and other features in two other motor phenotypes. Results did not differ after controlling for the effect of age (data not shown).

**Table 3.** The correlation (Spearman) between NLR and other characteristics in healthy controls and patients with Parkinson’s disease based on Tremor/PIGD score ratio.

|  |  | Age | Disease duration | Tremor score | PIGD score | UPDRS part III score | Modified Schwab & England ADL | UPSIT score | MoCA score | SCOPA-AUT score | SBR score in Caudate | SBR score in Putamen | SBR score in Caudate (ipsilateral) | SBR score in Putamen (ipsilateral) |
| --- | --- | --- | --- | --- | --- | --- | --- | --- | --- | --- | --- | --- | --- | --- |
| NLR, *P* value (rho) | **HCs (n = 148)** | 0.405 (0.069) | ... | ... | ... | 0.388 (-0.072) | ... | 0.133 (-0.124) | 0.704 (-0.031) | 0.480 (-0.059) | 0.590 (-0.045) | 0.702 (-0.032) | ... | ... |
|  | **All PD (n = 388)** | **0.011 (0.129)** | 0.495 (0.035) | 0.066 (0.093) | 0.662 (0.022) | 0.058 (0.096) | **0.015 (-0.123)** | 0.744 (-0.017) | 0.891 (-0.007) | 0.185 (0.067) | 0.199 (-0.066) | **0.026 (-0.114)** | **0.014 (-0.126)** | **0.002 (-0.158)** |
|  | **PIGD \* (n = 73)** | 0.991 (0.001) | 0.834 (0.025) | 0.706 (-0.045) | 0.667 (0.051) | 0.947 (-0.008) | 0.695 (0.047) | 0.732 (-0.041) | 0.232 (0.142) | 0.581 (0.066) | 0.947 (0.008) | 0.780 (-0.033) | 0.936 (0.010) | 0.942 (0.009) |
|  | **Indeterminate (n = 27)** | 0.813 (-0.048) | 0.488 (-.139) | 0.867 (0.034) | 0.921 (0.020) | 0.374 (-0.178) | 0.058 (-0.370) | 0.926 (0.019) | 0.524 (0.128) | 0.720 (0.072) | 0.97 (0.008) | 0.287 (-0.213) | 0.953 (0.012) | 0.659 (-0.089) |
|  | **TD (n = 288)** | **0.003 (0.172)** | 0.371 (0.053) | 0.050 (0.116) | 0.603 (0.031) | **0.009 (0.155)** | **0.015 (-0.144)** | 0.685 (-0.024) | 0.426 (-0.047) | 0.214 (0.073) | 0.106 (-0.096) | **0.041 (-0.120)** | **0.005 (-0.168)** | **<0.001 (-0.208)** |

Abbreviations: HCs, healthy controls; PD, Parkinson's disease; PiGD, Postural instability-gait difficulty; NLR, Neutrophil to lymphocyte ratio; UPDRS, Unified Parkinson's Disease Rating Scale; MoCA, Montreal cognitive assessment; Modified Schwab & England ADL, overall activities of daily living; UPSIT, University of Pennsylvania Smell Identification Test; SCOPA-AUT, Scales for Outcomes in Parkinson's disease; SBR, Striatal binding ratios.
\*The ratio of tremor and PiGD score in MDS-UPDRS. Parkinson's disease patients with a ratio of $\geq 1.15$ were classified as tremor-dominant (TD), ratio $\leq 0.90$ were classified as PiGD and $0.90 < \text{ratio} < 1.15$ were classified as intermediate.

Eventually, TD patients were tested for possible correlations between NLR and other characteristics, based on their H&Y stage. In 164 TD Parkinson’s disease patients at H&Y stage II, NLR had a positive correlation with age. As TD-patients in H&Y stage II were significantly older than TD patients in stage I (table 4), we carried partial correlation between NLR and SBR in putamen and caudate nuclei, controlling for age. Lower SBRs in bilateral putamen and caudate nuclei were significantly associated with higher NLRs in only the subgroup in stage II (table 5). No significant correlation was found in 124 TD patients with H&Y stage I.

**Table 4.** The correlation (Spearman) between NLR and other characteristics in patients with tremor dominant Parkinson’s disease (TD) based on Hoehn and Yahr stage.

|  |  |  |  |  |  |  |  |  |  |  | Contralateral* Nuclei |  | Ipsilateral Nuclei |  |
| --- | --- | --- | --- | --- | --- | --- | --- | --- | --- | --- | --- | --- | --- | --- |
|  |  | Age | Disease duration | Tremor score | PIGD score | UPDRS part III score | Modified Schwab & England ADL | UPSIT score | MoCA score | SCOPA-AUT score | SBR score in Caudate | SBR score in Putamen | SBR score in Caudate | SBR score in Putamen |
| NLR, *P* value (rho) | **H&Y stage I (n = 124)** | 0.902 (-0.011) | 0.548 (-0.055) | 0.563 (0.052) | 0.242 (-0.106) | 0.486 (0.063) | 0.064 (-0.167) | 0.263 (0.101) | 0.754 (0.028) | 0.980 (0.002) | 0.738 (-0.030) | 0.947 (-.006) | 0.640 (-0.043) | 0.541 (-0.056) |
|  | **H&Y stage II (n = 164)** | **0.002** (0.240) | 0.088 (0.134) | 0.140 (0.116) | 0.217 (0.097) | 0.278 (0.085) | 0.352 (-0.073) | 0.292 (-0.083) | 0.338 (-0.075) | 0.241 (0.092) | **0.040 (-0.162)** | **0.017 (-0.188)** | **0.003 (-0.221)** | **0.019 (-0.185)** |

Abbreviations: H&Y, Hoehn and Yahr stage; NLR, Neutrophil to lymphocyte ratio; UPDRS, Unified Parkinson's Disease Rating Scale; MoCA, Montreal cognitive assessment; Modified Schwab & England ADL, overall activities of daily living; UPSIT, University of Pennsylvania Smell Identification Test; SCOPA-AUT, Scales for Outcomes in Parkinson's disease; SBR, Striatal binding ratios.
\*Contralateral and ipsilateral nuclei are against the side of motor dominancy. Numbers for all SBRs are representative of added partial correlation between NLR and SBR in putamen and caudate nuclei, controlling for age.

**Table 7.** Demographic information and comparison of clinical outcomes in patients with Parkinson’s disease based on Hoehn and Yahr stage.

| Characteristic | H&Y stage I<br>(n = 124) | H&Y stage II<br>(n = 164) | P Value |
| --- | --- | --- | --- |
| Age, mean (SD), y | 59.0 (9.9) | 63.9 (8.9) | <0.001<sup>a</sup> |
| Disease duration, median (range), m | 4.0 [1-35] | 4.0 [0-36] | 0.864<sup>a</sup> |
| NLR, mean (SD) | 2.4 (0.9) | 2.7 (1.0) | 0.003<sup>a</sup> |
| SBR score in Caudate nucleus<br>(contralateral), mean (SD) | 1.9 (0.6) | 1.8 (0.5) | 0.166<sup>a</sup> |
| SBR score in Putamen nucleus<br>(contralateral), mean (SD) | 0.7 (0.3) | 0.7 (0.2) | 0.032<sup>a</sup> |
| SBR score in Caudate nucleus<br>(ipsilateral) , mean (SD) | 2.3 (0.6) | 2.1 (0.5) | 0.004<sup>a</sup> |
| SBR score in Putamen nucleus<br>(ipsilateral) , mean (SD) | 1.1 (0.4) | 0.9 (0.3) | <0.001<sup>a</sup> |
| Tremor score, mean (SD) | 0.5 (0.2) | 0.6 (0.3) | 0.017<sup>a</sup> |
| PIGD score, mean (SD) | 0.1 (0.1) | 0.2 (0.2) | 0.056<sup>a</sup> |
| MDS-UPDRS part III score,<br>mean (SD) | 15.1 (5.4) | 25.4 (8.5) | <0.001<sup>a</sup> |
| Modified Schwab & England ADL,<br>mean (SD) | 95.2 (5.1) | 92.0 (5.8) | <0.001<sup>a</sup> |
| MoCA score, mean (SD) | 27.3 (2.2) | 26.8 (2.4) | 0.074<sup>a</sup> |
| GDS score, mean (SD) | 2.1 (2.5) | 2.4 (2.3) | 0.081<sup>a</sup> |
| UPSIT score, mean (SD) | 23.7 (8.1) | 21.0 (8.0) | 0.006<sup>a</sup> |
| SCOPA-AUT score, mean (SD) | 9.0 (6.4) | 10.3 (6.1) | 0.051<sup>a</sup> |

Abbreviations: H&Y, Hoehn and Yahr stage; NLR, Neutrophil to lymphocyte ratio; SBR, Striatal binding ratios; PIGD, Postural instability-gait difficulty; MDS-UPDRS, Movement Disorder Society-sponsored revision of the Unified Parkinson's Disease Rating Scale; MOCA, Montreal cognitive assessment; GDS, Geriatric depression scale; Modified Schwab & England ADL, Overall activities of daily living; UPSIT, University of Pennsylvania Smell Identification Test; SCOPA-AUT, Scales for Outcomes in Parkinson's disease.
<sup>a</sup>Based on Mann-Whitney *U* test.

## 4. Discussion

In search of possible correlation between peripheral inflammation and striatal DAT availability in a large sample of drug-naïve early PD patients, we found that (1) NLR is significantly higher in PD patients than age and sex matched healthy controls. (2) Although NLR and striatal SBR did not differ between subtypes of PD in the early stages of the disease, only TD phenotype and more specifically at H&Y stage II confirmed predictive value of NLR to dopaminergic loss in bilateral striatal nuclei. (3) Worse motor severity was associated with higher NLR values in only TD phenotype.

Tremor is a cardinal motor feature of PD that has always exhibited unalike clinical course to other motor phenotypes. In contrast to TD, non-tremor dominant subtypes, especially PIGD predict worse and rapid disease progression with higher rates of non-motor features such as cognitive decline, mood disturbances, REM sleep behavior disorder and olfaction dysfunction (Jankovic and Kapadia, 2001; Burn et al., 2006; Kumru et al., 2007; Reijnders et al., 2009; Iijima et al., 2011; Konno et al., 2018). A longitudinal study has revealed that TD patients will not develop dementia until irreversibly convert to PIGD phenotype (Alves et al., 2006). Differed clinical pattern is also reflected on structural and functional assays. Autopsy of TD-PD patients has exposed less severe dopaminergic loss in SN than other PD subtypes (Paulus and Jellinger, 1991; Selikhova et al., 2013). This is in concordance with slower progression of symptoms in this group (Jankovic and Kapadia, 2001; Rajput et al., 2009), and is replicated on a recent probabilistic tractography study, which has revealed normal structural connectivity in nigro-pallidal and fronto-striatal circuits in TD patients, which was altered in non-TD (Barbagallo et al., 2017). Postmortem studies have also demonstrated distinct striatal pathology among different PD motor subtypes. While akinesia and rigidity evolve from cell loss in ventrolateral SNpc which project to the posterior putamen, degeneration in TD is most prominent in dorsomedial SNpc projecting to the caudate and anterior putamen, and to the retrorubral field with subsequent projections to the dorsolateral striatum and ventromedial thalamus (Jellinger, 1999). In support of this model, Eggers et al. showed lesions in caudate and lateral putamen (eagle-wing shape) on DATscans of TD patients, compared to lesions in dorsal putamen with egg-shape pattern of dopaminergic loss in akinetic-rigid phenotype (Eggers et al., 2011). This pattern is consistent with our finding of NLR correlation to lower dopamine uptake in both putamen and caudate in TD subgroup, while it was linked to lower DAT binding in putamen and not caudate in all PD patients. In fact, there is consistent report of higher extent of dopamine depletion in putamen than caudate in PD (Pikstra et al., 2016). Moreover, it is proposed that tremor severity may have a different mechanism than other motor features, as in contrast to bradykinesia and rigidity does not depend on the degree of reduced striatal dopamine uptake (Otsuka et al., 1996; Rossi et al., 2010; Eggers et al., 2012; Pikstra et al., 2016), but rather on relative contribution of reduced dopamine in putamen and caudate, indicative of more severe caudate involvement (Otsuka et al., 1996). Therefore, strong correlation of NLR to SBR in caudate of TD patients, but weak association with putamen agrees with these sightings. In this study, we did not find any difference between SBR values in ipsilateral or contralateral caudate and putamen nuclei between three examined subtypes with similar motor stage and severity. However, early drug-naïve TD patients of another study have shown higher DAT availability on bilateral putamen but no difference on caudate nuclei compared to akinetic-rigid PD patients with higher UPDRS-III and H&Y (Moccia et al., 2014). This may point to the possibly greater impact of putamen on motor severity in non-TD phenotype. In this study we did not investigate other neurotransmitters than dopamine which are conferred regarding tremorgenesis and its severity is shown to be mediated by reductions of serotonin transporter availability in the raphe nuclei more than that of dopamine in putamen (Qamhawi et al., 2015; Pasquini et al., 2018), or mediated by neurotransmitter imbalance between reduced GABAergic inhibitory influx from putamen confronted to intact dopaminergic input from SN to the internal pallidum (Rajput et al., 2008).

On the other hand, we found surprisingly, much more substantial reduction in DAT availability in ipsilateral putamen and caudate and a weak reduction in contralateral nuclei in TD patients at H&Y stage II compared to TD at stage I. This is also true for the strong correlation of NLR to ipsilateral caudate (p value= 0.003) and a weak relationship with contralateral caudate nucleus (p value= 0.04). Motor symptoms in PD are considered to evolve from dopaminergic depletion in contralateral nigrostriatal system. However, there are several recent reports of motor dominancy ipsilateral to the side of predominant dopaminergic deficit, especially in TD patients (Aguirregomozcorta et al., 2013; Erro et al., 2013; Hoshiyama et al., 2015). This is also true for medication naïve TD patients from PPMI cohort on early stages of the disease (Kaasinen, 2016). Another study on PPMI cohort (Pasquini et al., 2018) disclosed that a proportion of TD group at baseline progressed to bilateral limb involvement over 2 years follow up, which may be due to the observed ipsilateral striatal dopamine depletion in this study. It is of particular note that TD patients showed negative association between NLR and ipsilateral caudate, while splitting them based on H&Y stage, contralateral caudate emerged to have such association only in the group at stage II. More studies are needed to further elucidate the contribution of ipsilateral basal ganglia motor circuits to the tremorgenesis.

Outside the nigrostriatal tract, imaging studies have imposed the increased functional and metabolic activity of cerebellothalamocortical pathway in emerging parkinsonian tremor, which largely drives by depleted dopaminergic input from basal ganglia (Antonini et al., 1998; Helmich et al., 2011; Lewis et al., 2011). In a recent diffusion tensor imaging study, Wen et al. have revealed that drug-naïve early TD patients recruited from PPMI project have greater white matter integrity and lesser neural degeneration, while they observed widespread white matter dehiscence in PIGD group with similar motor stage and severity. The authors have discussed this finding as a neural compensation for nigrostriatal dopaminergic loss leading to more benign disease course in TD subtype. Additionally, White matter alterations were also more significantly related to severity of symptoms in PIGD than that of TD (Wen et al., 2018).

Overall, different pattern of clinical course and dopaminergic loss along with the consistent findings of involvement of distinct neural circuits outside the nigrostriatal region, strongly suggest the different underlying pathophysiology of PD tremor from that of other motor features (Thenganatt and Jankovic, 2014). This study is the first that has captured the different pathogenesis of the dopaminergic neuronal loss in terms of higher NLR interrelated to lower striatal dopamine in TD but not in non-TD Parkinson’s disease. NLR is an easy accessible marker of peripheral inflammation that is identified as a predictor of worse outcome in multiple chronic clinical and neurological disorders more prevalent in geriatric population, which also positively correlates to age in normal population (Li et al., 2015). Only two previous studies have investigated the association of NLR in idiopathic PD with opposite results. Akil et al. have found higher NLR in 51 PD (NLR= 3.1 ± 1.3) compared to 50 HC (NLR= 2.1 ± 10.3) (Akil et al., 2015), while Uçar et al. did not find any difference comparing 46 PD (NLR= 2.66 ± 1.05) and 60 HC (NLR= 2.46 ± 1.04) (Uçar et al., 2017). The later study also did not find any difference between TD and akinetic-rigid subtypes. These studies have assessed relatively small sample sizes of PD patients already on dopaminergic replacement therapy, which is shown to recover changes on T-lymphocytes (Kustrimovic et al., 2016). In this study we surveyed larger sample of 388 drug-naïve PD patients who showed higher levels of NLR (2.5 ± 0.9) compared to 148 age-matched healthy controls (NLR= 2.2 ± 0.8), although there was no such difference within PD subtypes. NLR is also shown to be attributed to lower connectivity in several white matter tracts implicated in early or prodromal PD pathology (Haghshomar et al., 2017).

Several studies have manifested that PD progression does not follow a linear model and decline in striatal uptake observed on DATscans are more rapid in early stages of the disease followed by a much more slower progression in advanced stages (Fearnley and Lees, 1991; Jennings et al., 2001; Nurmi et al., 2001). Therefore, it seems that early stages of the disease without the interference of dopaminergic replacement therapy, is the best point during the disease course to evaluate the contribution of inflammation to dopaminergic loss. A growing body of evidence suggests that presynaptic terminals in dorsal striatum undergo degeneration even prior to the SN and axonal degeneration should be the primary target for neuroprotective therapies (Tagliaferro and Burke, 2016). This is supported by in vivo DAT SPECT (Caminiti et al., 2017) and confirms that lower SBRs in putamen and caudate nuclei truly reflect the nigrostriatal pathology in early stages of PD. However, study on early stages impose some limitations that should be considered when interpreting our results. First, motor classification tends to change during the course of the disease, and our subtype classification was only based on screening visits. Second, tremor predominant PD patients recognize the symptoms and seek medical advice earlier and owing to the more benign course more readily cooperate with PPMI project that includes patients not receiving any parkinsonian treatment for 6 months. Therefore, higher number of cases tagged in TD subgroup and relatively fewer cases in PIGD and intermediate in PPMI cohort may have contributed to the false negative results in non-tremor predominant groups. More studies are clearly needed to investigate the generatability of these solid findings.

## Conclusion

In this research we investigated the association between immune and nervous system. PD patients were divided into three groups based on tremor/PIGD score, being TD, PIGD, and intermediate subgroups. We demonstrated that NLR can negatively predict SBR in bilateral putamen and caudate nuclei of TD group, and more specifically in stage 2 (H&Y) TD patients. Not observing such association in non-tremor predominant PD patients points to the different pathophysiology mechanisms between tremor and other motor symptoms in PD. Making a bridge between inflammation and neurodegeneration, it is suggested that peripheral inflammation can potentially contribute to the initiation and progression of PD, particularly the process of dopaminergic depletion in striatal regions. However, this relation between peripheral inflammation and parkinson progression is not conclusive. Much more studies are required to further investigate this subject.

## Ethical approval

All procedures performed here, including human participants were in accordance with the ethical standards of the institutional research committee and with the 1964 Helsinki declaration and its later amendments or comparable ethical standards.

## Informed consent

Informed consent was obtained from all individual participants included in the study.

## Consent for publication

All authors consent for the publication of this study.

## Availability of data and materials

All relevant data are available in the Parkinson’s Progression Markers initiative (PPMI) database (http://www.ppmi-info.org/data).

## Acknowledgements

Parkinsons Progression Markers Initiative (PPMI) database (www.ppmi-info.org/data) was our primary source of data in this research. For up-to-date information on the study, see www.ppmi-info.org. we gratefully thank all sponsors and funders of PPMI including the Michael J Fox Foundation for Parkinsons Research, AbbVie, Avid Radiopharmaceuticals, Biogen, Bristol-Myers Squibb, Covance, GE Healthcare, Genentech, GlaxoSmithKline (GSK), Eli Lilly and Company, Lundbeck, Merck, Meso Scale Discovery (MSD), Pfizer, Piramal Imaging, Roche, Servier, and UCB (www.ppmi-info.org/fundingpartners).

